# Large shifts in diatom and dinoflagellate biomass in the North Atlantic over six decades

**DOI:** 10.1101/2024.07.02.601152

**Authors:** Crispin M. Mutshinda, Zoe V. Finkel, Andrew J. Irwin

## Abstract

The North Atlantic Ocean has large seasonal blooms rich in diatoms and dinoflagellates which can contribute disproportionately relative to other primary producers to export production and transfer of resources up the food web. Here we analyze data from the Continuous Plankton Recorder to reconstruct variation in the diatom and dinoflagellate community biomass over 6 decades across the North Atlantic. We find: 1) diatom and dinoflagellate biomass has decreased up to 2% per year throughout the North Atlantic except in the eastern and western shelf regions, and 2) there has been a 1-2% per year increase in diatom biomass relative to total diatom and dinoflagellate biomass throughout the North Atlantic, except the Arctic province, from 1960-2017. Our results confirm the widely reported relationship where diatoms are displaced by dinoflagellates as waters warm on short time scales, but we did not observe a coherent effect of sea surface temperature.

## Introduction

North Atlantic phytoplankton communities are often dominated by blooms of diatoms in the spring and dinoflagellates in the late summer and early fall (Sverdrup 1953; Smayda 2002; Barton et al. 2013). Diatoms are characterized by their need for silicon for their cell wall, their tendency to form spring blooms in the mid and high latitudes, and are estimated to contribute approximately 40% of total marine primary production and particulate carbon exported to depth (Nelson et al. 1995; Jin et al. 2006; Tréguer et al. 2018). Dinoflagellates often bloom later than diatoms, are amongst the most genetically, morphologically, and trophically diverse groups of plankton; and include photosynthetic, mixotrophic and heterotrophic members (Pierella Karlusich et al. 2020).

Dinoflagellates are typically bi-flagellated with an organic cell wall formed from specialized vesicles (alveolae) that can form carbon-rich plates of various thicknesses (Bujak and Williams 1981; Janouškovec et al. 2017). Their C-rich walls, and often large genomes (Carnicer et al. 2023), can make their biomass higher in C:N and lower in N:P compared to the siliceous-walled diatoms (Menden-Deuer and Lessard 2000; Carnicer et al. 2021), influencing their food quality (Sterner and Elser 2002). In coastal waters, toxin production by some diatom and several dinoflagellate species can episodically disrupt fishing, aquaculture, and economic activities (Anderson et al. 2000; Hallegraeff 2003). Changes in the relative success of diatoms relative to dinoflagellates are expected to have a large impact on pelagic food web dynamics, fisheries, and the biochemical cycling of carbon, nitrogen, and phosphorus.

The Continuous Plankton Recorder (CPR) survey initiated by the Sir Alister Hardy Foundation for Ocean Science has been monitoring the plankton in the North Atlantic Ocean since the 1930s with millions of plankton abundance counts (Richardson et al. 2006; Batten et al. 2019). A silk mesh captures many larger phytoplankton species, predominantly diatoms and dinoflagellates. The methods used by the CPR survey have changed little since its initiation, providing an ideal dataset for quantifying how changes in physical and chemical conditions have altered plankton community structure and whether climate warming has left a detectable fingerprint on North Atlantic plankton community structure (Edwards et al. 2001; Richardson et al. 2006). Analyses of the CPR dataset suggest there have been shifts in the abundance of diatoms relative to dinoflagellates over time and with environmental conditions in some regions of the North Atlantic over the past half century – but that the direction of change may be regional. For example, Leterme et al. (2005) report increases in the CPR phytoplankton colour index (PCI), the abundance of dinoflagellates, and decreases in the abundance of diatoms in the North Atlantic over the period 1958-2002, especially over the Southeast Atlantic since 1988. In the central North Atlantic, Zhai et al. (2013) report a decadal increase in the ratio of dinoflagellates and a decrease in the ratio of diatoms, relative to all other phytoplankton, over 1985-2009. In contrast, Hinder et al. (2012) report an increase in the abundance of diatoms relative to dinoflagellates over 1960-2009, but with most of the increase occurring post 1995 in the northeast Atlantic (45-60°N, 15°W-10°E). Most recently, Holland et al. (2023) reported increases in diatom abundance in the CPR record (1960-2019) and many additional shorter near-shore time-series across the Greater North Sea but decreases in the Northeast Atlantic, in contrast dinoflagellate abundance showed decreases in the North East Atlantic and a mixture of increases and decreases in the Greater North Sea.

Changes in diatom and dinoflagellate community abundance over the last 60 years in the North Atlantic have been attributed to changes in temperature, water column stratification and nutrient supply, and wind-driven mixing (Leterme et al. 2005; Hinder et al. 2012; Hátún et al. 2017). It is difficult to determine if the variability in reported responses in diatom and dinoflagellates abundances across studies is due to differences in the physical regimes across the regions, the time periods analyzed, or differences in how the data were analyzed (Holland et al. 2023). Analyses across studies differ in the spatial and temporal subdivision of the North Atlantic examined and the taxa included in the analyses, potentially impacting conclusions. For example, Leterme et al. (2005) included species with a frequency of occurrence >1%, Hinder et al. (2012) examined the integrated abundance of diatoms and dinoflagellates present in >4% of samples and with persistent abundance across their timeframe of analyses, while Zhai et al. (2013) examined the total diatom and dinoflagellates abundance relative to all phytoplankton taxa identified. A major challenge in plankton time-series analysis is the large number of zero abundance data which is often overcome by focussing on dominant taxa and transforming data, both of which can induce biases (Mutshinda et al. 2022). Furthermore, interpretation of changes in abundance and the relative abundance of diatoms and dinoflagellates is complicated by large differences in the cell size and carbon content of the different diatoms and dinoflagellates making up the phytoplankton communities. We develop our analysis at the functional group level, rather than general or species, as many species vary neutrally within functional groups (Mutshinda et al. 2016), aggregations of species are inherently more predictable, and functional groups are a key target of modeling efforts and climate-change analysis. To more robustly assess if there have been temporal changes in diatom and dinoflagellates since 1960 across the North Atlantic, and if there are substantive differences in temporal trends across regions, we quantify the temporal trend in diatom and dinoflagellate biomass and a diatom index (= diatom/ (diatom + dinoflagellate) biomass) using a carbon content estimate of the species identified and a statistical model for the seasonal cycle, decadal-scale fluctuations, latitude, and the long-term trend (linear change over years) in 5 biogeographic provinces in the North Atlantic. Long-term trends in diatom and dinoflagellate biomass are well-constrained by the model, and are typically 1-2% per year, but vary in direction by province. The most robust trend is an average increase of 1-2% per year in the diatom index over the past 60 years across much of the North Atlantic, except for the Arctic where we observe a shift towards dinoflagellates at a comparable rate.

## Materials and Methods

### Data

Phytoplankton abundance data were obtained from the CPR project (Johns 2019). The CPR programme initiated by the Sir Alister Hardy Foundation for Ocean Science is the largest multi-decadal plankton-monitoring programme in the world (Richardson et al. 2006). The CPR sampling device is towed by ships of opportunity at their conventional operating speeds at a standard depth of 7 m. Seawater passes over a continuously moving band of silk mesh, which is stored in preservative for later analysis. Cells from approximately 3 m^3^ of seawater are collected for each sample, but the exact volume is unknown. CPR practice is to treat the data as abundances proportional to cell number density, but not to convert abundances to cells L^−1^ (Jonas et al. 2004; Richardson et al. 2006). The silk mesh captures many larger phytoplankton species, predominantly diatoms and dinoflagellates, as well as many smaller phytoplankton with diameters as small as 10 µm. Phytoplankton identified during zooplankton enumeration are sometimes recorded as present without quantification, but we discarded these presence-only data. For details, see the review by Richardson et al. (2006). Although phytoplankton data were first collected in the 1930s, broad spatial coverage of the North Atlantic, particularly in the Northwest North Atlantic, began in 1960. Each observation was associated with a particular Longhurst province (Longhurst 2007) using geographic boundaries provided by marineregions.org (http://marineregions.org/mrgid/22538, version 4, March 2010). We used only data from six Longhurst provinces: Atlantic Arctic (ARCT) and Boreal Polar (BPLR), North Atlantic Drift (NADR), Northeast Atlantic Shelves (NECS), Northwest Atlantic Shelves (NWCS), and Atlantic Subarctic (SARC). Data from ARCT and BPLR were combined into one region identified as ARCT due to data scarcity in these provinces separately (Fig. 1).

**Figure 1.**
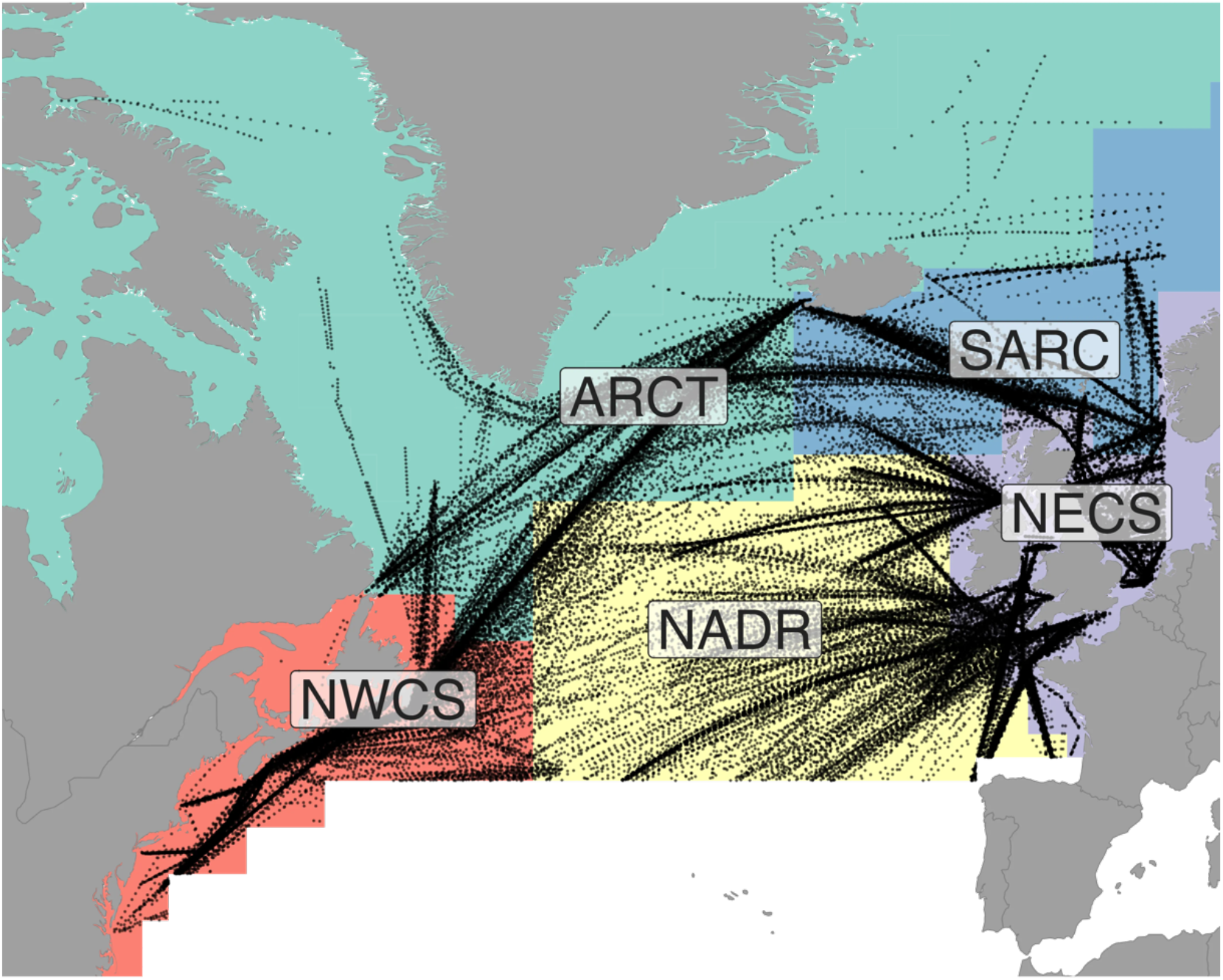
Map of five biogeographic regions and location of CPR samples used in this study. See Fig. S2 for sample size of diatom and dinoflagellate biomass data.

To calculate the community biomass for diatoms and dinoflagellates we summed the product of the CPR abundance for each taxon and the biomass for each taxon using an average cell volume for the taxon (Barton et al. 2013) and a cell volume to cell carbon model (Menden-Deuer and Lessard 2000). Sizes for taxa missing from this database were imputed as the mean log biomass for genus when known, or the mean log biomass for the functional type. Sea surface temperature (SST) is thought to be linked indirectly through changes in physical, chemical, and biological conditions to the absolute and relative prevalence of diatoms and dinoflagellates, but SST were not recorded simultaneously by the CPR project. We used the Hadley Centre sea surface temperature data product (HadISST, https://www.metoffice.gov.uk/hadobs/hadisst/) at 1° x 1° and monthly resolution (Rayner et al. 2003).

This region contains 393,975 observations of diatom and dinoflagellate species abundance with relatively few gaps in months or years (Table S1, Fig. S1). At the fine spatial resolution corresponding to a single square of sampling silk, it was common for a single sample (square of silk) to have 0 observations of diatoms or dinoflagellates, making it impossible to compute a biomass ratio at this scale. Data were aggregated to total functional group biomass within bins that were 2.5° wide in latitude, spanned the full longitudinal extent of the five provinces under consideration at monthly time resolution, yielding 12,162 observations of diatom biomass and 9,630 observations of dinoflagellate biomass (Tables S1, S2). Even after aggregation some locations had zero biomass of either diatoms (589 or 5%) or dinoflagellates (3121 or 24%). Almost half (45%) of the zero dinoflagellate biomass data were in the northernmost provinces (ARCT, SARC), consistent with our expectation of lower dinoflagellate biomass in colder waters (Table S3). Zero observations are a challenge for log-normally distributed data such as biomass since the log transformation drops out these zeros (Mutshinda et al. 2022). We computed the fraction of the combined diatom and dinoflagellate biomass due to diatoms, but zero biomass for either functional group was a problem here as we modelled the logit or log-odds of this biomass ratio. Zero biomass for both groups was replaced by 50% of the minimum community biomass over all samples in the aggregated dataset for each functional group (0.51 and 5.8 mg C, respectively for diatoms and dinoflagellates) and the distribution of the resulting data was examined to ensure it was approximately Normal. Our final aggregated data contained 12,751 observations of diatom and dinoflagellate biomass over five biogeographic provinces (observations per province varied from 1784 to 3837) and 6 decades (1960-2017, observations per decade ranged from 1491 to 2494). The diatom biomass, dinoflagellate biomass, and logit of the proportion of total (diatom + dinoflagellate) biomass due to diatoms, *h* = logit(*p*), were roughly Normally distributed for all biogeographic provinces under consideration (Fig. S2). Imputation of missing (i.e., zero) dinoflagellate biomass resulted in a spike in the biomass histogram which would likely have been dispersed over a larger range of biomass values if the observations had the resolution to resolve them. The histogram of the biomass ratio did not exhibit this spike because of the amount of variation in diatom biomass across samples with imputed dinoflagellate biomass. The speed of ships towing CPR samplers has changed over the years, somewhat affecting the volume of water collected per sample. As average ship speed has increased the volume of water filtered per sample has decreased, and thus we might anticipate a decrease in the apparent abundance. No significant correlation has been found between ship speed and the phytoplankton color index, which should be proportional to total phytoplankton biomass (Jonas et al. 2004). Our diatom index, being a ratio, should be insensitive to changing ship speed over time.

### Model specification

Our goal was to quantify the long-term linear rate of change in diatom biomass, dinoflagellate biomass, and the proportion of total (diatom + dinoflagellate) biomass due to diatoms, which we call the diatom biomass index. While the change in these quantities may be nonlinear in time, a linear approximation is a good first approximation to detect the difference between a directional trend in time and no trend. In addition to any long-term trend, we anticipated variation in diatom and dinoflagellate biomass and the diatom index arose within years with seasonal changes, across ocean biogeographic provinces, and with latitude within each province. The provinces subdivide the complexity of the North Atlantic, but each province spans a large region (Fig. 1, Table S3), and locations of some sampling routes have shifted within provinces over time. We used latitude and longitude as predictors during model development to account for spatial variation in sampling and oceanographic conditions within each province. Initial results (not shown) indicated that longitudinal variation was minimal, so we omitted it from our model. We modeled biomasses and the biomass index using linear regression with predictors corresponding to each of these sources of variation. The temporal trend may have been non-linear or year-to-year changes in environmental forcing may have resulted in fluctuations in the ratio around this trend. We considered two options for these deviations from a linear temporal trend: shifts in the ratio that changed each decade called the time-space model, or changes in the ratio correlated with sea-surface temperature (SST) anomaly relative to the mean predicted by other factors (month, year, province, and latitude) called the time-space-temperature model. Water temperature varies across provinces, with latitude, from month to month, and following a long-term trend. We used spatial and temporal variables to account for this variation in biomasses and the biomass index, so we did not include this variation in water temperature in our model. There was additional residual variation in temperature. For example, the temperature in a particular location and time is generally warmer (or cooler) than expected for that location, at that month, in that year. This additional variation is the temperature anomaly. It is uncorrelated with any of our spatial and temporal variables, so we included this temperature anomaly in a model to account for the effects of local variation in temperature on biomass and the diatom index.

We denoted the mean diatom and dinoflagellate biomass (µg C) by *q*_*A,l,m,y*_ and *r*_*A,l,m,y*_, respectively, in province *A* at latitude *l* during month *m* (January, February, …, December) of year *y*. The diatom index, equal to the proportion of total (diatom + dinoflagellate) biomass due to diatoms, is 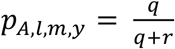. The bivariate datum (*p*, (1 − *p*)) is compositional as its components *p* and (1 − *p*) are bounded by 0 and 1 and sum up to 1 (Aitchison 1982). Bivariate compositional data are usually analyzed on the logit scale, where the logit transformation logit(*x*) = *l*n(*x*/(1 − *x*)) maps the interval (0,1) into the entire real line, allowing for the subsequent unconstrained data to be analyzed using conventional statistical techniques. Since compositional data only carry relative information, we also modeled spatio-temporal variation in each functional type’s biomass to gain insight into the magnitude and direction of biomass changes.

### Time-Space-Temperature models

Let *h*_*A,l,m,y*_ denote the logit-transformed diatom index in province *A* at latitude *l* during month *m* of year *y*. We modeled *h*_*A,l,m,y*_ as Normally distributed with mean

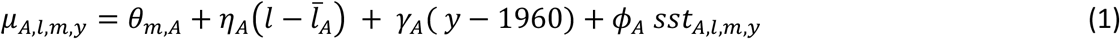

and variance 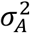, where *θ*_*m,A*_ is the effect of month *m, η*_*A*_ is the latitudinal gradient, with 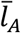 denoting the average latitude in province *A, γ*_*A*_ is the inter-annual trend, *sst*_*A,l,m,y*_ are sea-surface temperature anomalies, and *ϕ*_*A*_ is the linear effect of SST anomaly. We also modeled the ln-transformed biomass of each functional type and total biomass as in Eq. (1).

The SST anomaly was defined using a model for observed SST. Let *SST*_*A,l,m,y*_ denote the observed SST in province *A* at latitude *l* during month *m* of year *y*. We modeled this temperature as normally distributed with variance 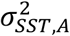 and expected value

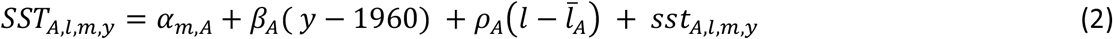

where *α*_*m,A*_, *β*_*A*_ and *ρ*_*A*_ the effect of month *m* (January through December), the inter-annual (year-to-year) trend, and the latitudinal gradient, respectively. The SST anomaly is the residual error after removing the effect of province, seasonal cycle, inter-annual trend, and latitude and was modeled as 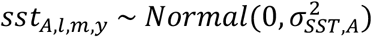. The SST anomaly in Eq. (1) is the posterior median of this quantity. We fit this model to SST data that coincided with the CPR sampling locations to develop the SST anomaly for the biomass and biomass-ratio models. We also fit this model to SST data sampled throughout the entire biogeographic province at 1° x 1° spatial resolution, except for the ARCT province which had large areas that were ice covered and extended far beyond the area surveyed by the CPR.

### Time-Space models

We modeled each of these quantities as Normally distributed with variance 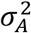 and expected value *µ* depending only on time and space by dropping the temperature anomaly from Eq. (1) and adding an effect for each decade:

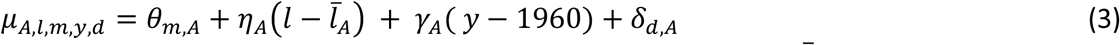

where *θ*_*m,A*_ is the effect of month *m, η*_*A*_ is the latitudinal gradient, with 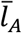 denoting the average latitude in province *A, γ*_*A*_ is the inter-annual trend, and *δ*_*d,A*_ is the effect of decade *d* (1960, 1970, …, 2010) for province *A*.

We completed the specification of our models with explicit statements of priors on all unknowns. For the parameters common to the Time-Space and Time-Space-Temperature models, we assumed exchangeable 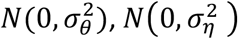, and 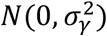 prior distributions on the month effects *θ*_*m,A*_, the inter-annual trends *γ*_*A*_, and the latitudinal gradients *η*_*A*_, with independent InvGamma(0.1,0.1) priors on the hyper-parameters 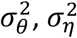, and 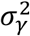, and placed Uniform (0,10) priors independently on the province-specific standard deviations *σ*_*A*_. On the temperature anomaly effects *ϕ*_*A*_ exclusive to the Time-Space-Temperature model, we placed exchangeable 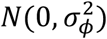 priors, with hyper-parameter 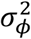 drawn from the InvGamma(0.1,0.1) distribution. For the temperature anomaly model we assumed exchangeable 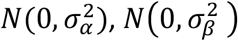, and 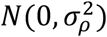 prior distributions on the month effects, the inter-annual trends, and the latitudinal gradients, with independent InvGamma(0.1,0.1) priors on the hyper-parameters 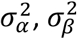, and 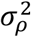, and placed Uniform (0,10) priors independently on the province-specific standard deviations *σ*_*SST,A*_. On the decadal effects *δ*_*d,A*_ exclusive to the Time-Space models, we assigned hierarchical priors implementing the Bayesian LASSO regularization (Tibshirani 1996; Park and Casella 2008), following the Extended Bayesian LASSO prior design (Mutshinda and Sillanpää 2010, 2012; Mutshinda et al. 2020), to selectively shrink the irrelevant effects toward zero. More specifically, we assumed that 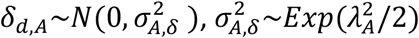, where *λ*_*A*_ = *λ ξ*_*A*_, *λ*∼Gamma(1,1) and *ξ*_*A*_ ∼Unif(0,2).

### Model fitting and validation

Since the joint posterior of the model parameters is not available in closed-form, we used Markov chain Monte-Carlo (MCMC) methods (Gilks et al. 1996), implemented in OpenBUGS (Thomas et al. 2006), to simulate from the posterior. We ran 10,000 iterations of three parallel Markov chains and discarded the first 4,000 iterations of each chain as burn-in period, thinning the remainder by a factor of 10. We assessed the convergence of the Markov chains informally through visual inspection of traceplots and autocorrelation plots. Model fits were assessed using posterior predictive checks (see Supplementary Methods). The importance of predictor variables was assessed by omitting one predictor at a time and observing the decrease in the proportion of variance in the response explained by the model (Bayesian *R*^2^). We used posterior predictive model checking to validate our models, which is the standard approach with Bayesian modelling (Gelman et al. 2013). Our posterior predicive model checking compared observed data with data simulated from the model using histograms and summarized differences using Bayesian p-values (see Supplementary Methods).

## Results

Our results revealed an increasing long-term trend in the diatom index (diatom biomass divided by total (diatom + dinoflagellate) biomass) in four of the five North Atlantic biogeographic provinces under consideration (NADR, NECS, NWCS, and SARC), with yearly average increases of 0.7-2.5% in NADR, NWCS, SARC, and NECS. By contrast, the long-term trend in the proportion of total biomass due to diatoms is negative in the ARCT province with a yearly average decrease of 0.9% (Fig. 2, Table S4). There was a long-term decrease in diatom biomass of 0.7-0.9% in ARCT and SARC, the two northernmost provinces and a long-term increase of 1.8-2.2% per year in NWCS and NECS. For dinoflagellate biomass, the long-term trend is decreasing in SARC, NADR and NECS with average rates of decrease 0.2-2.2%, whereas the trend in NWCS is increasing at a yearly rate of 1.2%. The average rate of decrease for total (diatom + dinoflagellate) biomass was 0.2-1.4% in the SARC, NADR, and ARCT, respectively with average rates of increase of 0.6-1.1% in NECS and NWCS. An annual increase of 1 or 2% per year compounds to an increase of 80% and 230% over 60 years, and a decrease of 1 or 2% per year compounds to a decrease of 45% and 70% over 60 years.

**Figure 2.**
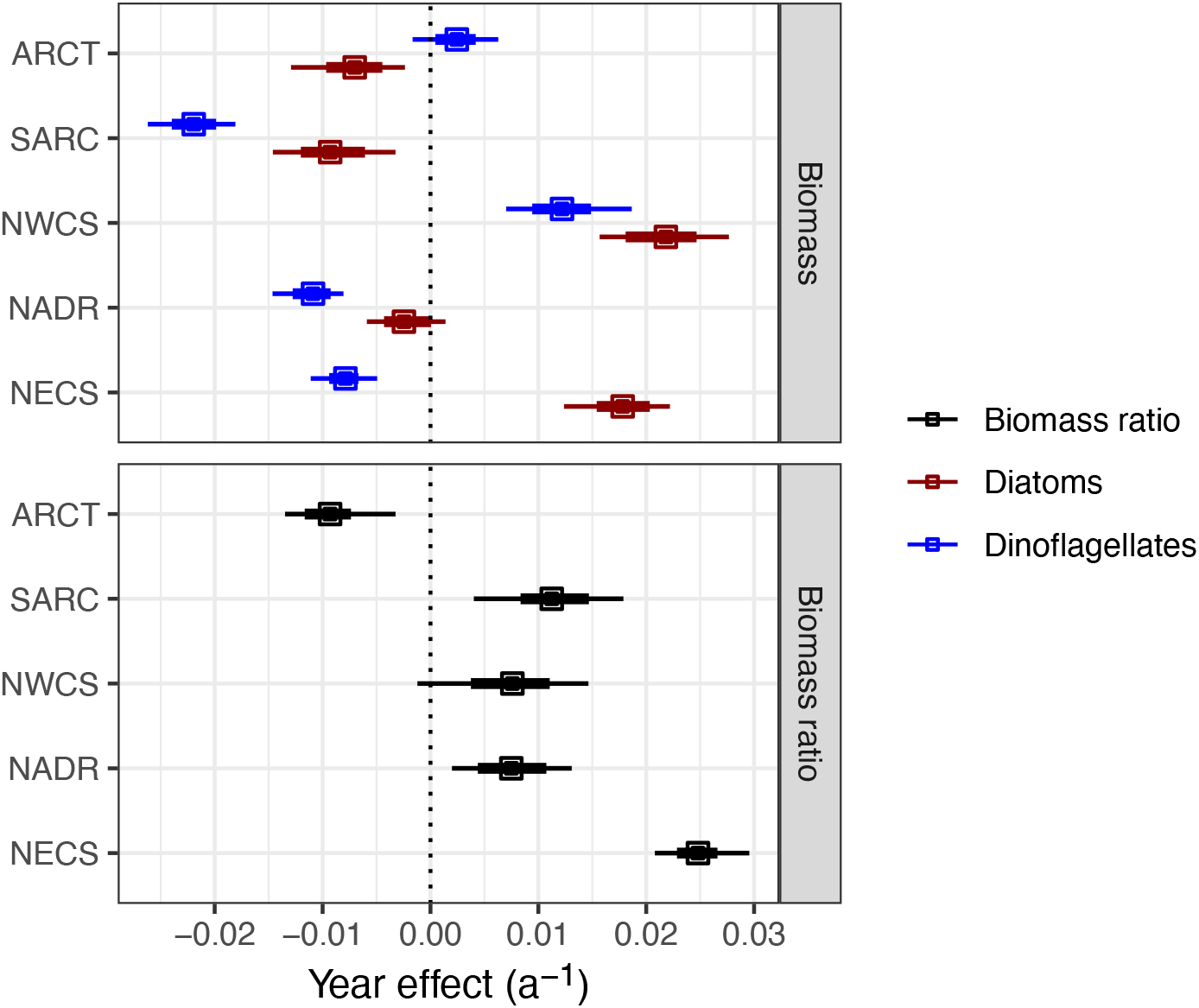
Average annual rate of change in biomass (top panel: diatoms, red; dinoflagellates blue; log scale) and the biomass ratio (bottom panel: diatom biomass / (diatom + dinoflagellate biomass), logit scale) in the five biogeographic provinces estimated from a time, space and temperature model. An effect (slope) of 0.01 a^−1^ corresponds to approximately a 1% change in biomass or the biomass ratio per year. Vertical dashed lines emphasize no change. Points are the median of the posterior distribution and error bars are 95% (thin) and 66% (thick) credible intervals. See Fig. S3 for a comparison between two models.

In all biogeographic provinces, the intra-annual variation in the relative biomass of diatoms vs. dinoflagellates was characterized by strong seasonal cycles with a peak between March and May, and minimum value in August (Fig. 3, bottom). In all five provinces, the intra-annual cycles in diatom biomass displayed a peak between April and June coinciding with the spring bloom. Following this peak, the diatom biomass declined as the dinoflagellate biomass increased toward its climax occurring in July (Fig. 3, top). In NWCS, the dinoflagellate biomass exhibited the lowest intra-annual variability, remaining relatively high and continually high throughout the year, consistent with reports of relatively high dinoflagellate biomass in the NWCS, particularly in the Winter (Johns et al. 2003a). The annual mean biomass is higher for dinoflagellates compared to diatoms in all provinces except the ARCT (horizontal lines, Fig. 3, Table S5). Dinoflagellate relative to diatom biomass was highest in NWCS. Dinoflagellate biomass was lower than diatom biomass, on average, only in the Arctic, consistent with this province currently being a less favourable habitat for dinoflagellates and greater potential for environmental change to shift the Arctic community towards dinoflagellates due to long-term changes in stratification and warming. Total diatom + dinoflagellate biomass was highest in NECS and lowest in the ARCT.

**Figure 3.**
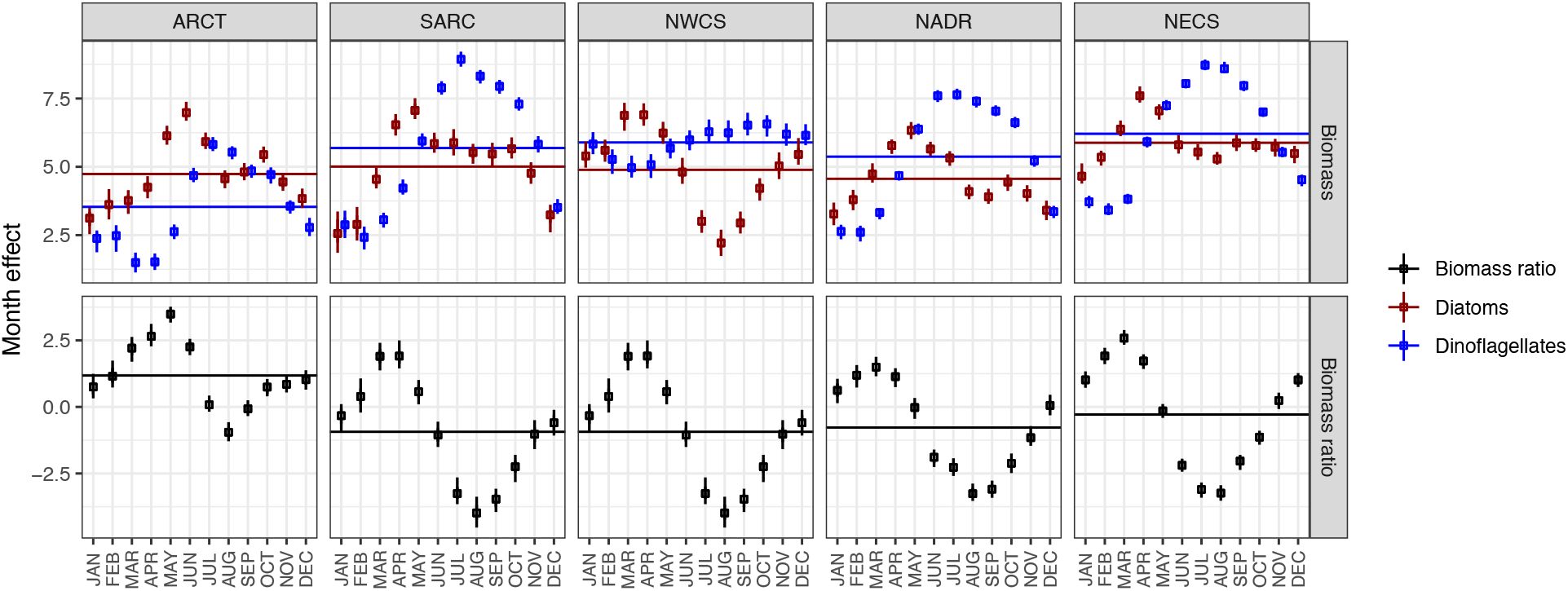
Mean monthly biomass (top panels: diatoms, red; dinoflagellates, blue; log scale) and biomass ratios (bottom panels: diatom biomass / (diatom + dinoflagellate biomass); logit scale) for five biogeographic provinces estimated from a time, space and temperature model. Vertical dashed lines emphasize 0 biomass (top) or equal biomass in diatoms and dinoflagellates (bottom). Points are the median of the posterior distribution and error bars are 95% (thin) and 66% (thick) credible intervals. See Fig. S4 for a comparison between two models.

We computed an SST anomaly from the Hadley SST data product and a model of SST that accounted for variation by month, over years, across provinces, and with latitude within each province. The SST anomaly was used as a predictor for the diatom index and phytoplankton biomasses. In all provinces, the SST anomaly correlated negatively with the diatom index (Fig. 4, bottom right), indicating anomalously warmer SST favored higher dinoflagellate relative to diatom biomass. The SST anomaly correlated positively with the biomasses of both functional types, indicating enhanced net phytoplankton biomass at warmer SST, but the dinoflagellate response was relatively larger (Fig. 4, top right). The effect of SST anomaly on diatom and dinoflagellate biomass in the NWCS province was nearly equal, resulting in almost no effect on the diatom index. The biomass and diatom index trends with latitude were mixed (Fig. 4, left side). In the SARC and NADR provinces, diatoms were relatively more dominant in the north, while in the NWCS the community shifted towards dinoflagellates towards the north.

**Figure 4.**
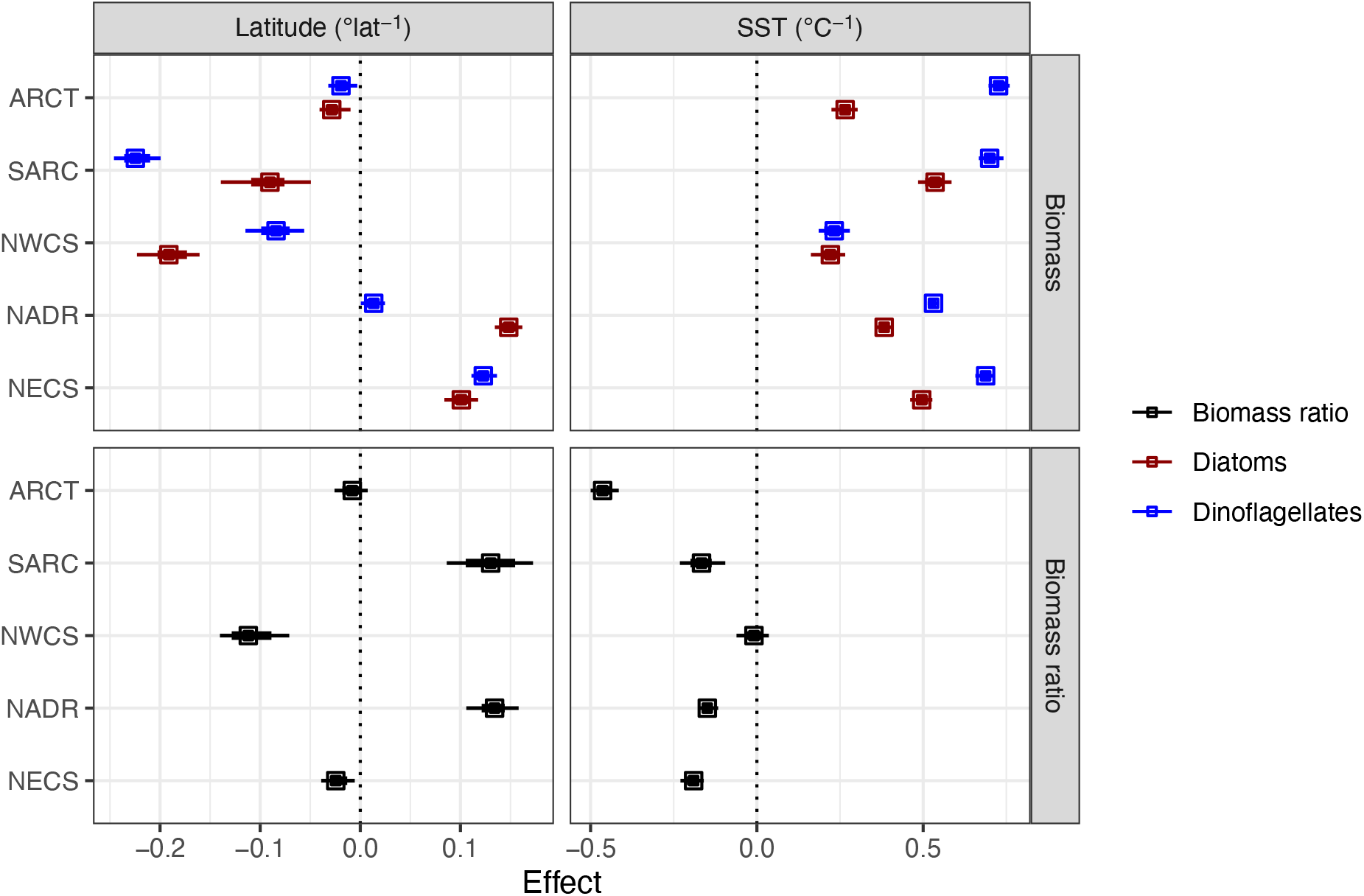
Mean effect of latitude (left panels) and temperature anomaly (right panels) for biomass (top panels: diatoms, red; dinoflagellates, blue; log scale) and biomass ratios (bottom panels: diatom biomass / (diatom + dinoflagellate biomass); logit scale) for five biogeographic provinces estimated from a time, space and temperature model. An effect (slope) of 0.1 corresponds to approximately a 10% change in biomass or the biomass ratio per °latitude or °C. Vertical dashed lines emphasize 0 change. See Fig. S5 for a comparison between two models.

The posterior means for the effects of month, year, province, and latitude from our main model (Time-Space-Temperature) were essentially identical to the results of the Time-Space models. The Time-Space models lacked the SST anomaly predictor and contained a decadal effect omitted in the Time-Space-Temperature model. The SST anomaly had predictive power and reduced uncertainty in these estimated parameters (Figs. S3, S4, S5). Since the SST anomaly was constructed to be independent of the other predictors (month, year, province, latitude), it did not affect the median values of the corresponding parameters. The uncertainty (95% credible interval) in the posterior means was generally smaller for the Time-Space-Temperature model, most notably for the estimates of the inter-annual trends on biomass and biomass ratio (Fig. S2). The posterior distributions of the decadal effects from the Time-Space models were clustered around zero and the 95% credible intervals generally included 0 (Fig. S6), implying that the nonlinear temporal changes in the diatom index and phytoplankton biomasses were negligible. Some of the effects were large (±0.5, corresponding to a change of 40-60% relative to the mean) but with a correspondingly large uncertainty. These results support the use of a linear trend over years, since the data did not reveal a non-linear trend on the decade scale.

We anticipated the possibility of a non-linear change in the diatom index and biomasses with time, which we modelled as step-wise changes for each decade. While we found some evidence for non-linear changes on the decade scale, the effects were highly variable from decade to decade and our estimates were very uncertain (Fig. S6). We found that the temperature in a patch of water relative to the mean temperature as predicted by location, month, and year (the temperature anomaly) had a notable effect on biomass and the diatom index (Fig. 4) and reduced the posterior uncertainty of the effects of month and year as well (Fig. S2, S3).

While temperature change over time (year and month) was not a primary focus in this work, we quantified and removed this variation to compute a temperature anomaly and so we report the results of that analysis. The SST data exhibited the expected seasonal cycles (Fig. S7) with similar results at the CPR sampling locations (blue) and province-wide results (black). In the NWCS, the CPR sampling locations were generally cooler by 2-3°C than the province as a whole and in the other provinces there were notable differences in the temperature at CPR sampling locations and throughout the province in the winter months (except for the ARCT province for which this comparison was not made.) The latitudinal trend in temperature was negative as expected (cooler in the north of each province) and similar whether the CPR sampling locations or whole province was studied (Fig. S8, left). Sea-surface temperatures exhibited an increasing trend of about 0.01-0.02°C a^−1^ across the provinces, except at the CPR sampling locations for SARC and NADR which showed no increase in temperature and a decrease in temperature over the study period, respectively (Fig. S8, right). The Hadley SST data show a statistically significant linear increasing trend of similar magnitude when studied at 1° x 1° spatial resolution throughout almost the entire study region (Fig. S9).

Variable importance for the Time-Space-Temperature model assessed from the changes in the Bayesian *R*^2^ by omitting the predictors month, latitude, and temperature anomaly one at a time showed that month accounted for most of the variance followed by temperature anomaly. Latitude had only a minor explanatory effect (1-2% of total variance, Table S6). The total variance explained by the Time-Space-Temperature model ranged from 32-64%. Model checking showed that the posterior distribution of the means and variances of the diatom index and diatom and dinoflagellate biomass from the Time-Space-Temperature models well approximated the data (Figs. S10, S11).

## Discussion

There is accumulating evidence that climate warming over the last century is altering ocean conditions, primary production, chlorophyll concentration, and the biogeography of the key phytoplankton taxa (Levitus et al. 2005; Behrenfeld et al. 2006; Boyce et al. 2010). Diatoms and dinoflagellates are two of the major phytoplankton functional groups in the North Atlantic and changes in their relative biomass may impact export production (Tréguer et al. 2018). It has been hypothesized that climate warming will shift plankton communities towards dinoflagellates and away from diatoms, altering food web composition with potentially significant consequences for fisheries (Leterme et al. 2005; Xiao et al. 2018). The CPR data have been used to investigate trends in the biomass of phytoplankton, diatom and dinoflagellate functional groups, and the relative abundances of diatoms and dinoflagellates on many occasions, but a lack of standardization in how the plankton community has been assessed across regions (e.g., subset of species examined, statistical framework, using sums of abundances vs. biomass) and temporal range analyzed makes rigorous North Atlantic regional-temporal comparisons challenging. Here we synthesize nearly 60 years of data (1960-2017) from the Continuous Plankton Recorder project spanning much of the northern North Atlantic divided into 5 provincial regions (Fig. 1) to identify if there has been a long-term trend in diatom and dinoflagellate biomass and the proportion of their biomass contributed by diatoms (diatom index) and whether these changes vary regionally. In contrast to expectations, we find there has been temporal increase in the diatom index over much of the North Atlantic, except the Arctic region, and the sum of the total biomass of the diatoms and dinoflagellates increases in the shelf regions but decreases in the central and sub-Arctic regions.

Our analysis confirms that there are stark regional differences in how diatom and dinoflagellate biomass has changed in the North Atlantic since the 1960s (Fig. 2). There were large increases in diatom biomass in the eastern and western shelf regions (NWCS, NECS), diatom biomass was essentially unchanged in the central North Atlantic (NADR), and decreased in the polar regions (SARC, ARCT). Dinoflagellate biomass increased in the NWCS, was unchanged in the ARCT, and decreased in the other three provinces. The decline in diatoms in the NADR, sub-Arctic and Arctic are results are broadly consistent with a global model of ocean biogeochemistry coupled to a climate model (Bopp et al. 2005). There is relatively little comparable observational data from the Arctic and North Atlantic Drift province, but Zhai et al. (2013) found a decrease in diatom relative to diatom + dinoflagellate abundance in the ARCT and NADR over 1991-2009, consistent with our longer-term trend. While Zhai et al. (2013) link these changes in community structure to the NAO, sea surface temperature and stratification, Hátún et al. (2017) hypothesize changes in the subpolar gyre and decreases in dissolved silicate may be responsible. Also consistent with our observations Osman et al. (2019) found primary productivity and diatom and dinoflagellate abundance has been in decline in the sub-Arctic, Head and Sameoto (2007) reported increases in both diatom and dinoflagellate abundance between 1962-1971 versus 2001-2003 on the Newfoundland shelf, eastern Scotian shelf and central/western Scotian shelf, and Johns et al. (2003b) found that dinoflagellate abundance has been increasing in the Grand Banks region in winter. There is a wealth of studies from the Northeast Coastal Shelf, particularly in the North Sea, but reported results are not unambiguous. For example, Leterme et al. (2005) found a decline in diatom abundance in the NE and SE Atlantic especially since the late 1960s/early 1970s, Hinder et al. (2012) found an increase in diatom relative to dinoflagellate abundance using a subset of diatoms species related to sea surface temperature and windy conditions, while Edwards et al. (2022) found a north-south contrast, with an increase in diatom abundance with warming in the more northerly versus more southerly part of the region. It is difficult to compare the results across these studies and establish the extent of regional differences across the North Atlantic without a consistent analysis of the entire dataset.

Our analysis identified distinct regional differences in the long-term temporal trend in diatom and dinoflagellate biomass by accounting for intra-annual, interannual, and spatial variability in the diatom index and diatom and dinoflagellate biomass. Our analysis captured the strong interannual trend, including expected differences in timing of peak biomass of both functional groups (diatoms before dinoflagellates), and delay in blooms further north (Fig. 3). We assessed, alternatively, a decadal variation to allow for a non-linear trend with time and a temperature anomaly, finding that the temperature anomaly approach produced the most explanatory power and reduced uncertainty in the parameter estimates (Fig. S3, S4, S5). Spatial variation was incorporated using the standardized Longhurst provinces and latitude to allow for variation within provinces, while still permitting the spatial aggregation necessary to observe a strong signal. The regular CPR routes have sometimes shifted northward or southward over time, leading to additional variation within provinces that can be partially attributed to the latitude of the observation (see Edwards et al. (2022) for a recent detailed description of the CPR dataset).

Regional differences in the response of diatom and dinoflagellate abundance have been correlated to climate indices such as the NAO and associated differences in hydroclimatic conditions such as wind, mixed layer depth and upper water column stratification, that in turn affect nutrients and conditions such as sea surface temperature and light, e.g., (Leterme et al. 2005; Hinder et al. 2012; Edwards et al. 2022). Interactions between environmental conditions and trophic transfer to zooplankton and fish may also be altering the timing and abundance of diatoms and dinoflagellate populations, through differential effects of temperature on phytoplankton versus zooplankton or trophic cascades (Edwards and Richardson 2004; Frank et al. 2005; Rose and Caron 2007). Given there has been warming throughout most of the North Atlantic over the past 6 decades (Polyakov et al. 2010; Levitus et al. 2012; Bates and Johnson 2020)(Fig. S9) and a diversity of responses in diatom and dinoflagellate biomass across regions, there is clearly no universal link between the NAO or increases in sea surface temperature and the temporal trajectory of diatom or dinoflagellate biomass.

Although there is no coherent response of diatom or dinoflagellate biomass with increasing sea surface temperatures in the North Atlantic since the 1960s, anomalous temperature increases, after accounting for province, latitude, month, and year, result in a lower diatom index (shift towards dinoflagellates) in all provinces except the NWCS (Fig. 4). We speculate that the NWCS community is already relatively dinoflagellate dominated, limiting the potential for further shifts towards dinoflagellates (Fig. 3). This almost universal response may be a reflection of a direct effect of increasing temperature on growth rates, leading to increased biomass. Indirect effects of temperature can be positive or negative through correlated increases in stratification and windiness, affecting turbulence and the availability of light and nutrients. Differential effects of temperature on grazers and their prey may alter food web structure, diatom and dinoflagellate biomass and the diatom index. Increases in temperature anomaly shift the community towards dinoflagellates, while the long-term trend with time (and increasing temperature) shifts the community towards diatoms in most regions. We hypothesize that these short-term variations in temperature are too rapid for diatoms and dinoflagellates to adapt and therefore more likely to result in eco-physiological change in community structure. Independent analyses in other parts of the ocean and experimental evolution studies in the lab show that phytoplankton can adapt to changes in temperature over several tens to hundreds of generations (Irwin et al. 2015; Schmidt 2016; Ajani et al. 2018; Benner et al. 2020). Our inference is that plankton adapt evolutionarily to temperature change and to predict or interpret the effects of decades of temperature change requires a deeper analysis than a simple effect of long-term warming of the ocean.

Here we document a long-term shift towards diatoms and away from dinoflagellates, on a biomass basis, throughout the temperate and sub-polar North Atlantic. This shift towards diatoms was the result of both increasing and decreasing trends in diatom and dinoflagellate biomass, depending on the biogeographic province. This analysis reveals a signature of a large-scale restructuring of the North Atlantic phytoplankton community not typically anticipated as the consequences of climate change. To document this trend required decades of investment in consistent sampling. Data from the CPR plankton survey are as valuable as they are rare. Continued investment in these surveys is essential for developing an understanding of the ocean-climate system. Although we reported results for the Arctic province, we recognize a severe under-sampling of the spatial extent of this province and caution our results are only representative of the southernmost extent of the Arctic and Boreal province in the North Atlantic. In the other four provinces, we see good agreement in the rate of warming at CPR sampling sites and the whole province in the NWCS and NECS, but note that the average temperature at the sites sampled by the CPR in the SARC and NADR decreased over 6 decades, despite the average temperature in these provinces increasing (Figs. S8, S9). The hallmark of our analysis is that the trends are highly variable across provinces, reflecting the consequences of regional changes in climate (e.g., increased windiness in NECS) and differences in the baseline community composition across provinces.

Notably, dinoflagellate biomass was much lower in the Arctic resulting in a higher diatom index and dinoflagellate biomass was only weakly seasonal in the NWCS (Fig. 3), so it seems reasonable to anticipate distinctive responses to climate change in these provinces.

Assuming the underlying mechanisms do not change, the next few decades could bring further decreases in diatom and dinoflagellate biomass, with a shift towards diatoms in much of the North Atlantic and a shift towards dinoflagellates in the Arctic. These changes have likely had notable consequences for carbon export and the amount of biomass transferred up the food web. Future climate change might have similar effects, although non-linear or unanticipated changes are certainly possible. Extrapolating the long-term effects of climate change from this analysis is challenging because of the opposing factors documented by the long-term change (Fig. 2) and the temperature anomaly (Fig. 4). The effect of a 1°C temperature anomaly on diatom and dinoflagellate biomass and the diatom index is approximately equal, but opposite, to several decades of long-term climatic change (Figs. 2, 4). Changes to future carbon export and trophic transfer will depend on the relative importance of long-term trends and short-term variation arising from temperature change, sea-ice melt, changes in freshwater inputs and circulation.

## Data availability

All data are publicly available in repositories and cited in the references.

## Acknowledgments

This work was supported by grants from the Simons Foundation (549935 to AJI, 549937 and 986772 to ZVF), the Ocean Frontier Institute (NWABCP to AJI and ZVF), and Discovery grant awards from the National Science and Engineering Research Council of Canada (AJI, ZVF).

We thank the CPR survey team past and present. The CPR survey is supported by funding from Canada, France, Ireland, The Netherlands, Portugal, the UK and the USA. The survey depends on the voluntary co-operation of owners, masters and crews of merchant vessels which tow the CPRs on regular routes. HadISST data were obtained from https://www.metoffice.gov.uk/hadobs/hadisst/ and are British Crown Copyright, Met Office provided under a Non-Commercial Government Licence http://www.nationalarchives.gov.uk/doc/non-commercial-government-licence/version/2/. The authors declare that they have no conflicts of interest.

## Notes

### Competing Interest Statement

The authors have declared no competing interest.

