## Supplement for "Large shifts in diatom and dinoflagellate biomass in the North Atlantic over six decades"

Supplementary Materials for

### Long-term changes in diatom and dinoflagellate biomass in the North Atlantic highlight regional consequences of climate change

Crispin M. Mutshinda[1], Zoe V. Finkel[2], Andrew J. Irwin[1]

[1]Department of Mathematics and Statistics, Dalhousie University, Halifax, NS, Canada

[2]Department of Oceanography, Dalhousie University, Halifax, NS, Canada

**Contents**

Supplementary Methods

Supplementary Results

Supplementary Tables S1-S6

Supplementary Figures S1-S11

#### Supplementary Methods

##### *Model assessment using posterior predictive checks*

The standard approach to Bayesian model validation is posterior predictive model checking (Gelman et al. 1996; 2013), which draws on the ability to approximate the posterior predictive distribution, i.e., the distribution of unobserved values conditional on observed data. The posterior predictive distribution of an observable $\tilde{y}$ conditional on observed data $y$ is defined by

$$p(\tilde{y} \mid y) = \int L(\tilde{y} \mid \boldsymbol{\theta})\, p(\boldsymbol{\theta} \mid y) d\boldsymbol{\theta} \quad \text{(S1)}$$

where $L(y|\boldsymbol{\theta})$ and $p(\boldsymbol{\theta})$ are respectively the likelihood and the prior in the Bayesian model. The posterior predictive distribution is esssentially the likelihood of $\tilde{y}$ integrated over the posterior uncertainty in $\boldsymbol{\theta}$. The optimal Bayesian prediction under a quadratic loss function is the posterior predictive mean value $E[\tilde{y}|y]$.

The rationale of posterior predictive model checking is to compare the observed data with replicated data simulated from the posterior predictive distribution, or alternatively to compare some test quantity $T(y, \boldsymbol{\theta})$ based on the observed data, to the same statistic $T(y^{rep}, \boldsymbol{\theta})$ for replicated data from the posterior predictive distribution. If the model fits the data well, then replicated data generated under the model should look similar to the observed data. Therefore, systematic discrepancies between $T(y, \boldsymbol{\theta})$ and $(y^{rep}, \boldsymbol{\theta})$ indicate model misfit. The comparison of $T(y, \boldsymbol{\theta})$ to $(y^{rep}, \boldsymbol{\theta})$ may be visual using graphical tools such as histograms, or formal using posterior predictive $p$-values also known as Bayesian $p$-values defined as (Gelman et al. 1996; 20013)

$$P_B = Pr(T(y^{rep}, \theta) \geq T(y, \theta|y) \quad \text{(S2)}$$

If observed data are consistent with model predictions, then $P_B$ should be close to 0.50. Values of $P_B$ close to 0 or 1 provide evidence for model inadequacy with $P_B$ close to 0 indicating a lack of fit and values close to 1 pointing to overfitting, which arises when a model is unnecessarily too complex. In the Bayesian MCMC framework, a simulation-based approximation of $P_B$ is given by the proportion of posterior predictive data replicates for which the test quantity exceeds its original data counterpart. At the most basic level, posterior predictive checks involve the comparison of observed data to their posterior predictions. Relevant test quantities can differ depending on the assumed sampling distribution. For the normal likelihood assumed here to describe the logit of the proportion of total (diatom + dinoflagellate) biomass due to diatoms and the log biomass of each functional type, the relevant test statistics are the mean and the standard deviation, which fully characterize the normal distribution.

#### Supplementary Results

*Counts of non-zero observations of diatom and dinoflagellate biomass in the CPR data at different spatial resolutions*

The raw CPR data analysed here had 304,472 observations of diatoms and 217,232 observations of dinoflagellates, but most of these observations did not have simultaneous counts for both diatoms and dinoflagellates. When aggregated to 1° x 1° monthly resolution, the number of diatom and dinoflagellate biomass observations per month ranged from 2,400 to 40,000 (Table S1). In about half the year, the number of diatom biomass observations was double (or more) the number of dinoflagellate biomass observations, resulting in poor estimates of the biomass ratio at this resolution. When the aggregation was increased to 2.5° latitude, five provinces, and monthly the total number of observations per province and month dropped considerably (Table S2) but the number of diatom and dinoflagellate observations were much more balanced. The number of observations of diatom and dinoflagellate biomass at this aggregation level varied within years and across decades, with the smallest number of observations arising in the winter months. A notable feature was the near-total absence of data for the 1980s in the NWCS (Fig. S10).

#### Supplementary Tables

**Table S1.** The number of observations of diatom and dinoflagellate biomass for the CPR data aggregated by month at 1° x 1° spatial resolution.

| | JAN | FEB | MAR | APR | MAY | JUNE | JUL | AUG | SEPT | OCT | NOV | DEC |
|---|---|---|---|---|---|---|---|---|---|---|---|---|
| **DIATOM** | 8167 | 11952 | 28418 | 46906 | 48554 | 34292 | 24138 | 19989 | 24283 | 27376 | 19119 | 11278 |
| **DINOFL.** | 3208 | 2446 | 4305 | 9776 | 20971 | 30857 | 39379 | 36999 | 30621 | 22762 | 10681 | 5227 |

**Table S2.** The number of observations of diatom (upper number) and dinoflagellate (lower number) biomass for the CPR data aggregated by month over 2.5° latitude bands for each biogeographic province.

| | JAN | FEB | MAR | APR | MAY | JUNE | JUL | AUG | SEPT | OCT | NOV | DEC |
|---|---|---|---|---|---|---|---|---|---|---|---|---|
| **ARCT** | 94 | 65 | 103 | 162 | 229 | 251 | 253 | 242 | 254 | 220 | 208 | 162 |
| | 30 | 24 | 15 | 24 | 88 | 174 | 227 | 216 | 189 | 160 | 121 | 69 |
| **SARC** | 36 | 62 | 128 | 168 | 191 | 185 | 172 | 163 | 173 | 168 | 156 | 97 |
| | 14 | 15 | 46 | 93 | 144 | 178 | 180 | 173 | 180 | 160 | 130 | 52 |
| **NWCS** | 139 | 133 | 146 | 167 | 165 | 156 | 132 | 111 | 133 | 147 | 151 | 154 |
| | 130 | 111 | 111 | 123 | 133 | 139 | 143 | 147 | 157 | 149 | 137 | 140 |
| **NADR** | 115 | 164 | 230 | 299 | 287 | 273 | 256 | 236 | 250 | 240 | 218 | 172 |
| | 39 | 51 | 105 | 194 | 240 | 274 | 264 | 263 | 266 | 236 | 188 | 92 |
| **NECS** | 221 | 268 | 330 | 347 | 349 | 338 | 344 | 337 | 326 | 317 | 305 | 264 |
| | 114 | 127 | 180 | 288 | 334 | 335 | 345 | 343 | 325 | 299 | 235 | 171 |

**Table S3.** Summary of missing observations after spatial aggregation into 2.5° latitude bins. Number of biomass ratios, diatom biomasses (observed/missing), dinoflagellate biomasses (observed/missing), the latitude range of observation locations, and the number of missing temperature data in spatially aggregated data, grouped by biogeographic province.

| Province | Biomass ratio (n) | Diatoms (n / missing) | Dinoflagellates (n / missing) | Latitude range | SST (# missing) |
|---|---|---|---|---|---|
| ARCT | 2324 | 2243 / 81 | 1337 / 987 | (50, 80) | 0 |
| SARC | 1784 | 1699 / 85 | 1365 / 419 | (57.5, 72.5) | 0 |
| NWCS | 1859 | 1734 / 125 | 1620 / 239 | (30, 52.5) | 2 |
| NADR | 2947 | 2740 / 207 | 2212 / 735 | (42.5, 57.5) | 0 |
| NECS | 3837 | 3746 / 91 | 3096 / 741 | (45, 65) | 0 |

**Table S4.** Average annual rate of change in the diatom index (diatom/(diatom + dinoflagellate) biomass ratio, logit scale) and in the biomass of each functional type and the total of both groups (natural log scale) across the five biogeographic provinces estimated from the Time-Space-Temperature models, expressed as a % change per year ± half the width of the 95% credible interval. An effect (slope) of 0.01 $a^{-1}$ corresponds to approximately a 1% change in biomass or in the biomass ratio per year.

| | **ARCT** | **SARC** | **NWCS** | **NADR** | **NECS** |
|---|---|---|---|---|---|
| Diatom index | -0.935 ± 0.51 | 1.1 ± 0.69 | 0.76 ± 0.79 | 0.75 ± 0.55 | 2.5 ± 0.44 |
| Diatom biomass | -0.7 ± 0.53 | -0.93 ± 0.57 | 2.2 ± 0.6 | -0.245 ± 0.36 | 1.8 ± 0.49 |
| Dinoflagellate biomass | 0.245 ± 0.4 | -2.2 ± 0.41 | 1.2 ± 0.58 | -1.1 ± 0.33 | -0.79 ± 0.31 |
| Total biomass | -0.16 ± 0.4 | -1.4 ± 0.36 | 1.1 ± 0.49 | -0.59 ± 0.33 | 0.575 ± 0.33 |

**Table S5.** Posterior median of the annual mean diatom index (dimensionless) and phytoplankton biomass (diatom, dinoflagellate, total) (g C) ± half the width of the 95% credible interval.

| | **ARCT** | **SARC** | **NWCS** | **NADR** | **NECS** |
|---|---|---|---|---|---|
| Diatom index | 1.2 ± 0.17 | -0.93 ± 0.3 | -0.93 ± 0.3 | -0.77 ± 0.19 | -0.28 ± 0.15 |
| Diatom biomass | 4.7 ± 0.16 | 5 ± 0.21 | 4.9 ± 0.24 | 4.6 ± 0.13 | 5.9 ± 0.17 |
| Dinoflagellate biomass | 3.5 ± 0.16 | 5.7 ± 0.14 | 5.9 ± 0.22 | 5.4 ± 0.11 | 6.2 ± 0.1 |
| Total biomass | 5.5 ± 0.16 | 6.8 ± 0.14 | 7.2 ± 0.16 | 6.4 ± 0.11 | 7.5 ± 0.11 |

**Table S6.** Proportion of variation in the data explained by Time-Space-Temperature models excluding each explanatory variable one at a time.

| | Model | | |
|---|---|---|---|
| | Diatom index (Logit) | Biomass | |
| Excluded variable | | Diatom | Dinoflagellate |
| None (Full model) | 0.37 | 0.32 | 0.64 |
| SST | 0.34 | 0.22 | 0.41 |
| Latitude | 0.36 | 0.30 | 0.62 |
| Month | 0.06 | - | - |

#### Supplementary Figures

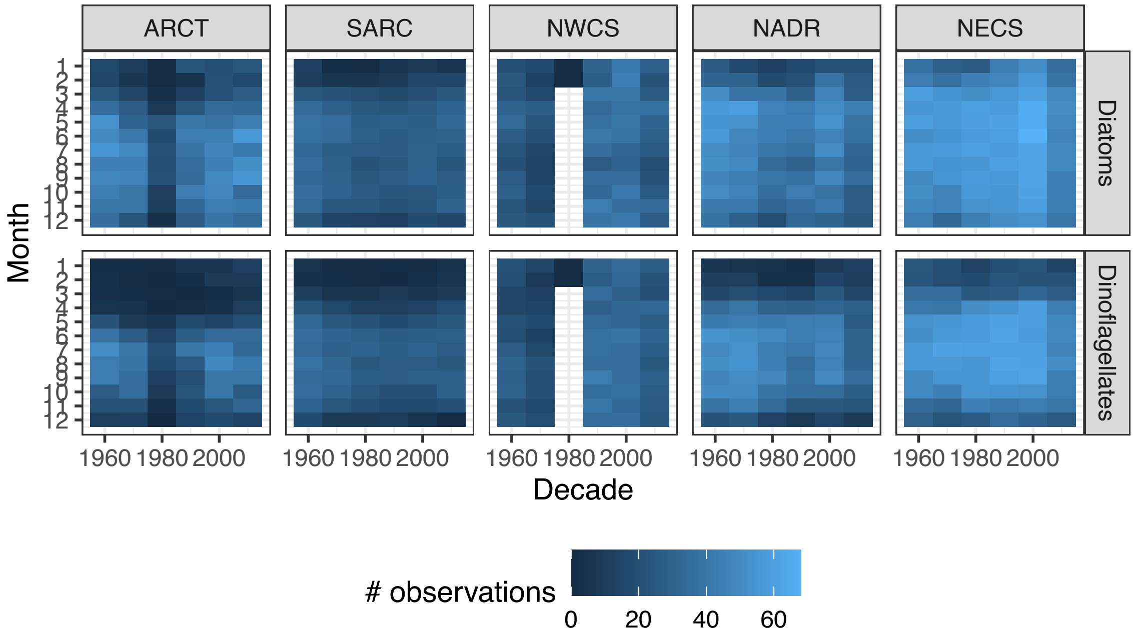


**Figure S1.**

Number of observations of diatom (top panels) or dinoflagellate (bottom panels) biomass after aggregation into 2.5° latitude bins in each province by month and decade.

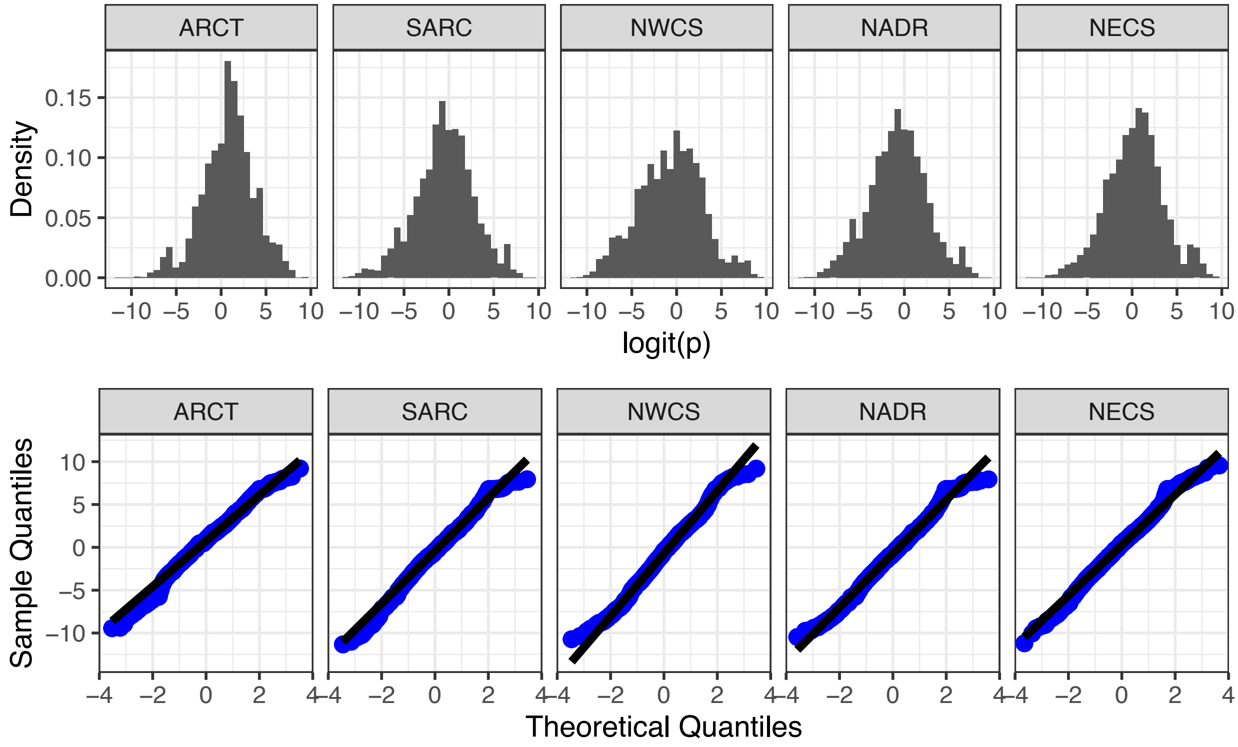


**Figure S2.**

Histograms (top row) and quantile-quantile plots (bottom row) of the logit of the diatom index (diatom biomass / (diatom + dinoflagellate biomass)) for each province showing that its distribution is approximately Normal. There are spikes at 0 and 1 in the untransformed ratios due to imputation of missing values, but these are not noticeable after the logit transform.

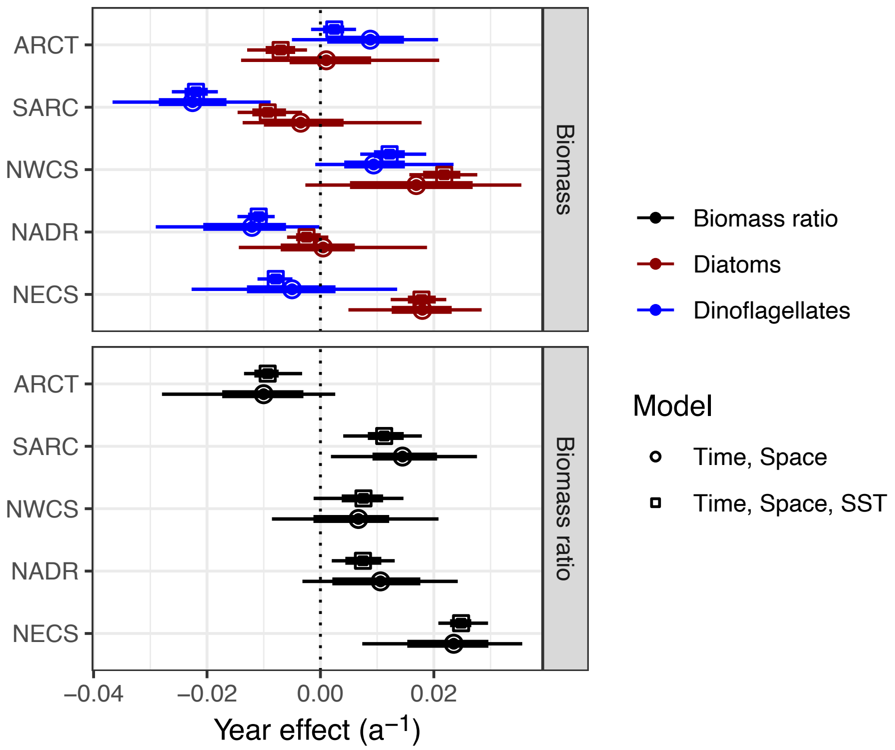


**Figure S3.**

Average annual rate of change in biomass (top panel: diatoms, red; dinoflagellates blue, log scale) and the diatom index (bottom panel: diatom biomass / (diatom + dinoflagellate biomass); logit scale) in the five biogeographic provinces estimated from two models (time, space and temperature model, squares, upper points, also shown in Fig. 2; time and space, circles, lower points). An effect (slope) of 0.01 $a^{-1}$ corresponds to approximately a 1% change in biomass or the biomass ratio per year. Points are the median of the posterior distribution and error bars are 95% (thin) and 66% (thick) credible intervals. See Fig. 3 for a simplified plot with one model.

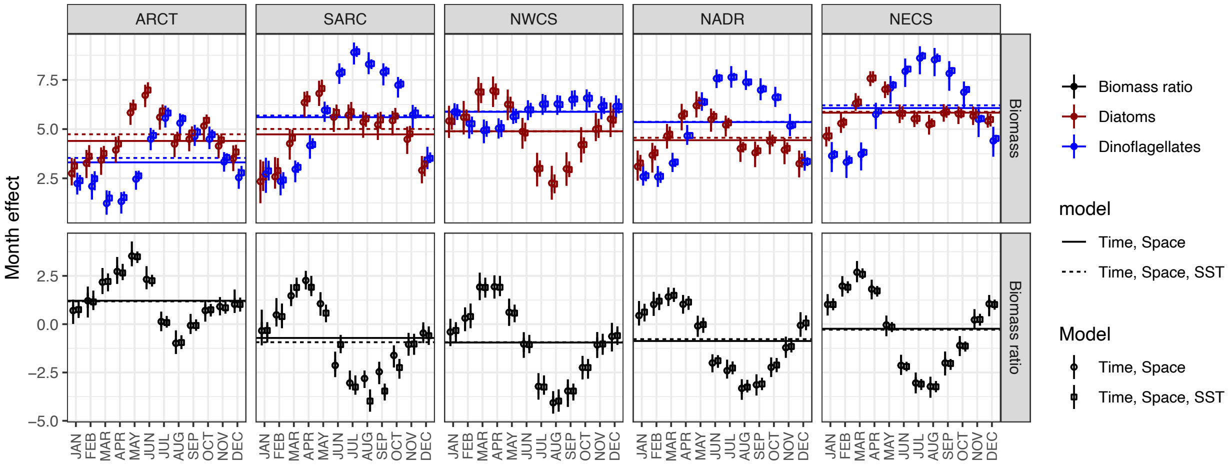


**Figure S4.** Mean monthly biomass (top panels: diatoms, red; dinoflagellates, blue; log scale) and diatom index (bottom panels: diatom biomass / (diatom + dinoflagellate biomass); logit scale) for five biogeographic provinces estimated from two models (Time-Space, circles, left points; time-space-SST model, squares, right points). Points are the median of the posterior distribution and error bars are 95% (thin) and 66% (thick) credible intervals. See Fig. 2 for a simplified plot with one model.

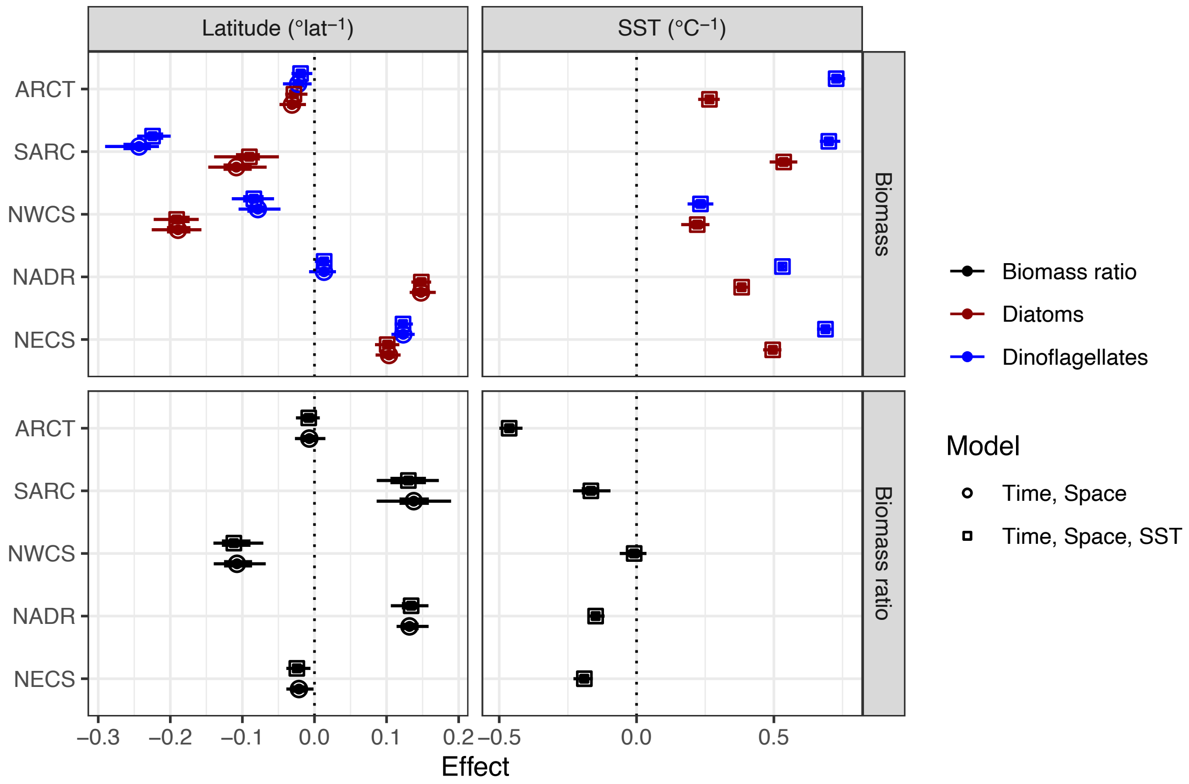


**Figure S5.** Mean effect of latitude (left panels) and temperature anomaly (right panels) for biomass (top panels: diatoms, red; dinoflagellates, blue; log scale) and biomass ratios (bottom panels: diatom biomass / (diatom + dinoflagellate biomass); logit scale) for five biogeographic provinces estimated from two models (time and space, circles, left points; time, space and temperature model, squares, right points). An effect (slope) of 0.1 corresponds to approximately a 10% change in biomass or the biomass ratio per °latitude or °C. Vertical dashed lines emphasize 0 change. Points are the median of the posterior distribution and error bars are 95% (thin) and 66% (thick) credible intervals. See Fig. 4 for a simplified plot with one model.

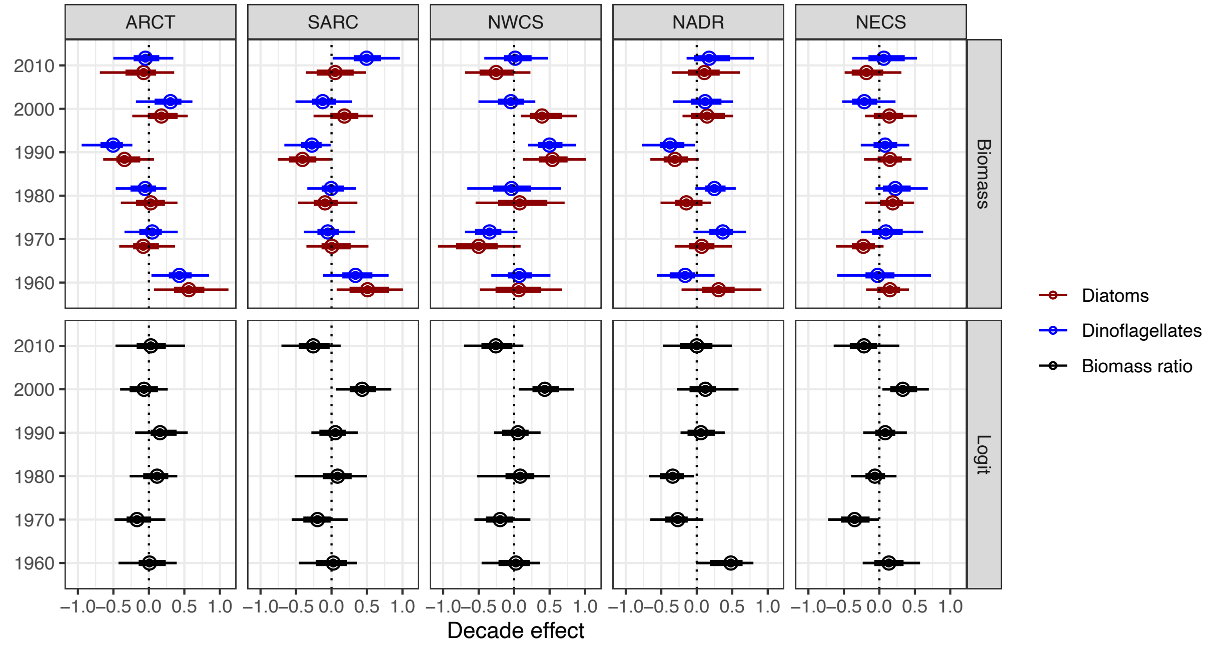


**Figure S6.** Variation in mean biomass (top panels: diatoms, red; dinoflagellates, blue; log scale) and biomass ratio (bottom panels: diatom biomass / (diatom + dinoflagellate biomass); logit scale) for five biogeographic provinces across decades estimated from the Time-Space model. Points are the median of the posterior distribution and error bars are 95% (thin) and 66% (thick) credible intervals. Vertical dashed lines emphasize 0 change.

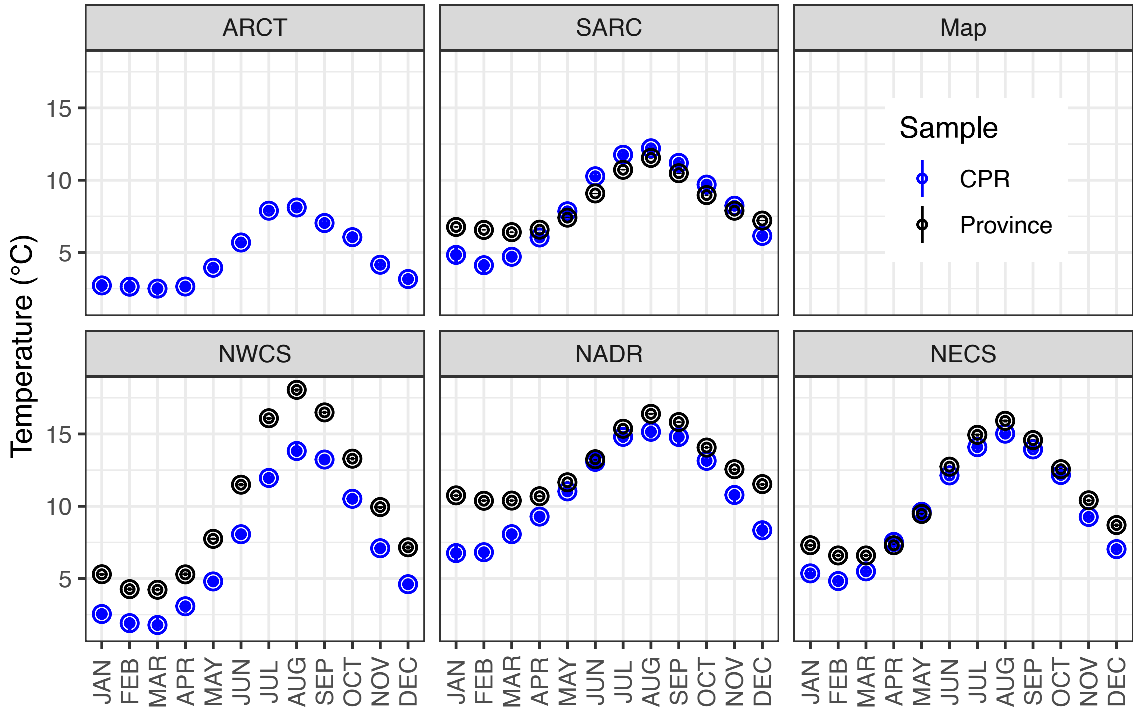


**Figure S7.** Mean monthly temperature (°C) in each biogeographic province estimated from Hadley sea surface temperature reanalysis product sampled at locations where CPR data observed (blue) and throughout each province at 1° resolution (black). Points are the median of the posterior distribution and error bars are 95% (thin) and 66% (thick) credible intervals. Error bars are all smaller than symbols. Province-wide sampling was not done for the ARCT province as its spatial extent is much larger than the CPR sampling and much of the region is at -1.8°C.

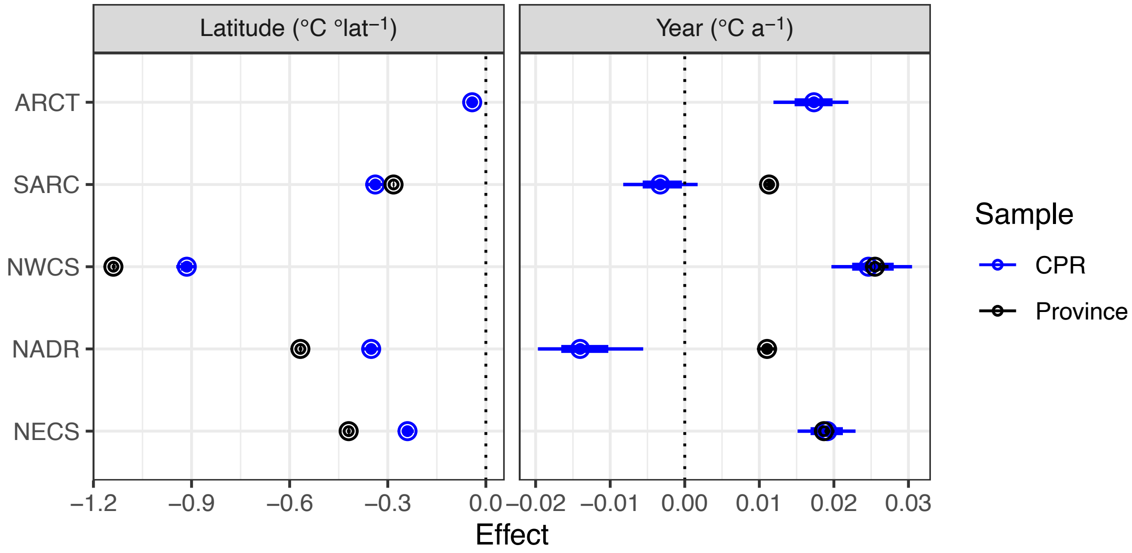


**Figure S8.** Change in mean temperature as a function of latitude (left panel, °C °lat$^{-1}$) and year (right panel, °C a$^{-1}$) in each biogeographic province estimated from Hadley sea surface temperature reanalysis product sampled at locations where CPR data observed (blue) and throughout each province at 1° resolution (black). Points are the median of the posterior distribution and error bars are 95% (thin) and 66% (thick) credible intervals. Error bars are all smaller than symbols for the latitude estimate. Vertical dashed lines emphasize 0 change. Province-wide sampling was not done for the ARCT province as its spatial extent is much larger than the CPR sampling and much of the region is at -1.8°C.

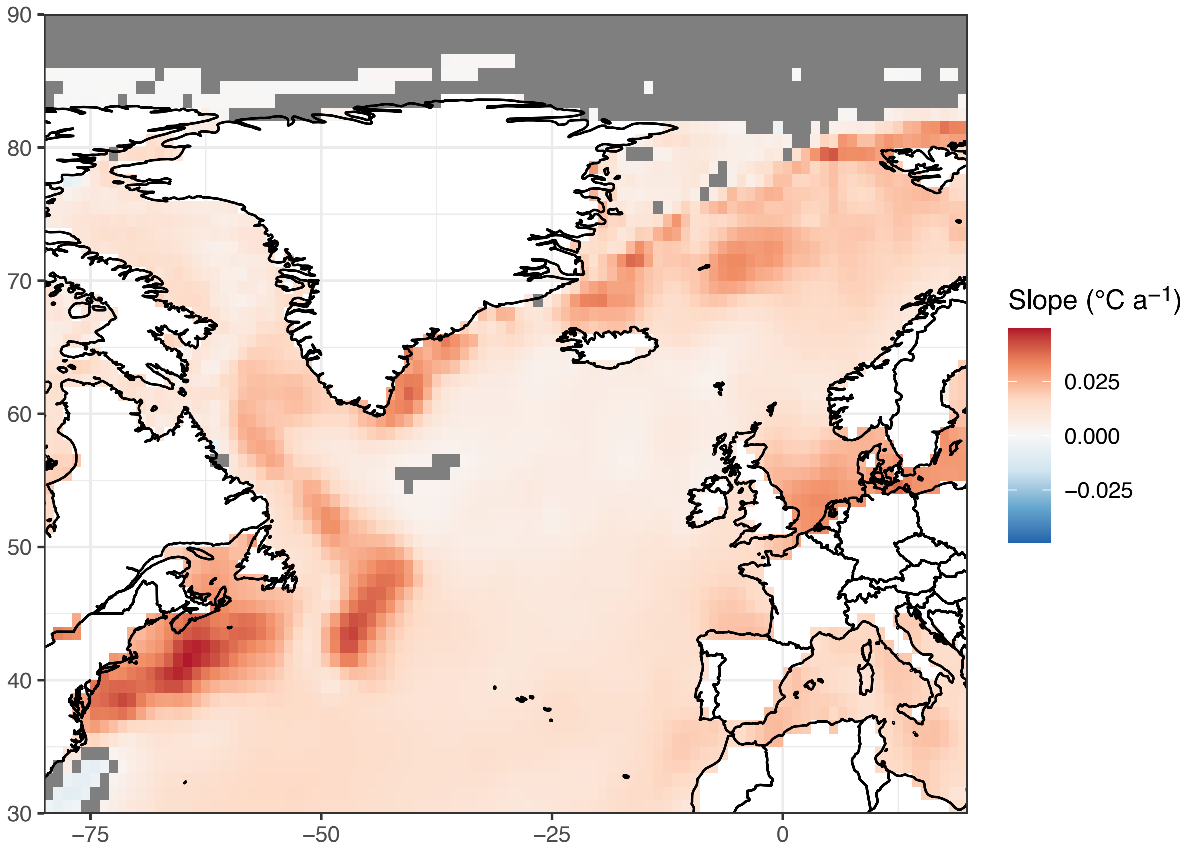


**Figure S9.** Mean rate of sea surface temperature change (°C $a^{-1}$) from monthly Hadley SST over the period 1960-2017 on a 1° grid. Slopes were estimated by linear regression. Non-significant slopes ($p > 0.05$) masked with gray boxes.

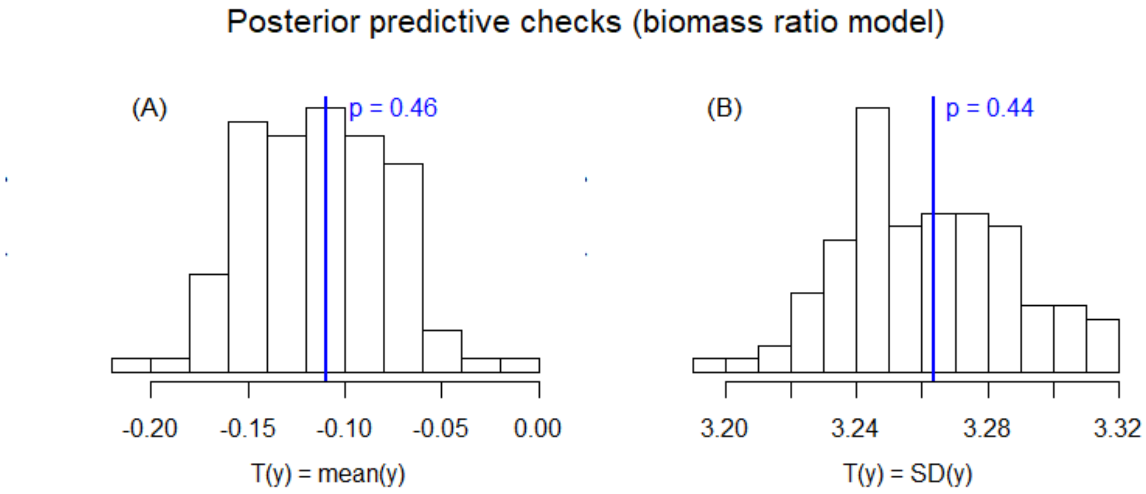


**Figure S10.** Histograms of posterior predictive means (left) and variances (right) over 100 posterior predictive of replicates for the logit of the proportion of total (diatom + dinoflagellate) biomass due to diatoms under the Time-Space-SST logit model. The vertical line represents the observed value of the test statistic. The Bayesian p-values displayed in each panel are all in the range 0.25-0.75, far from the extremes 0 and 1, implying that the model predictions do not systematically deviate from the data.

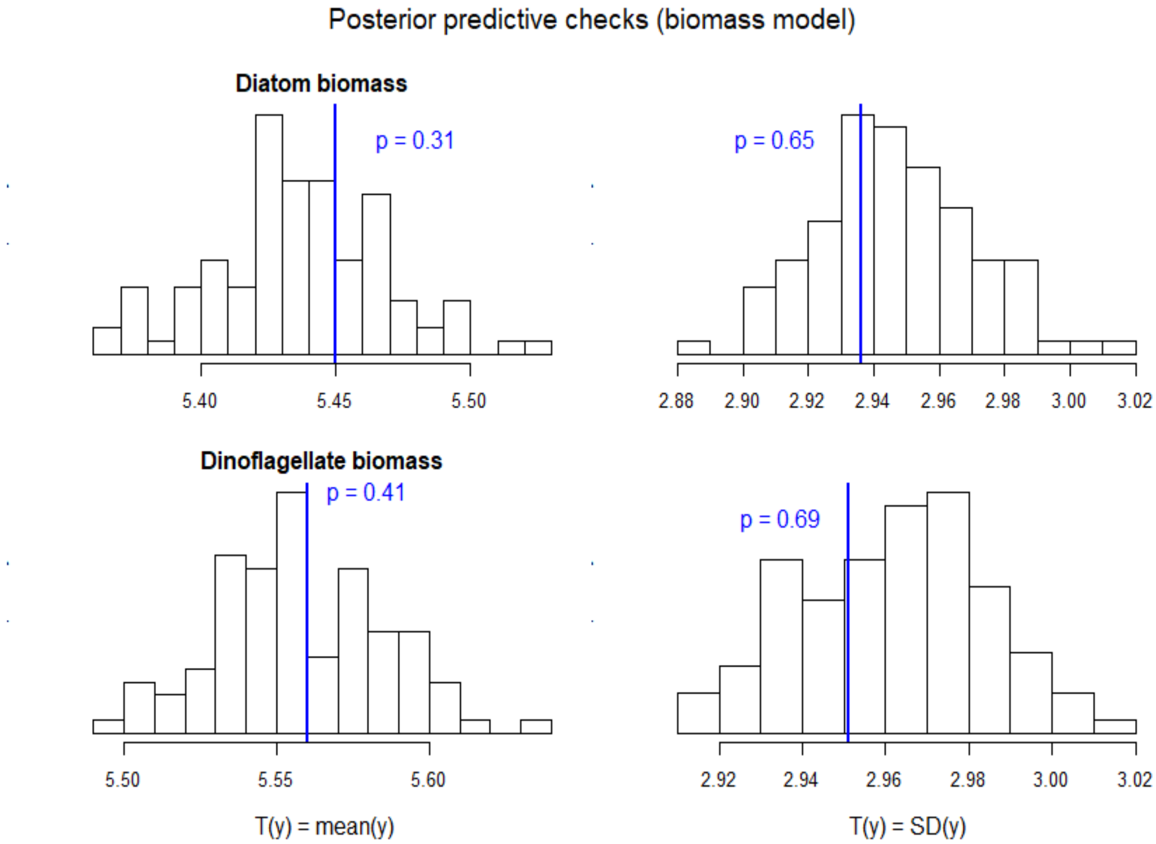


**Figure S11.** Histograms of posterior predictive means (left) and variances (right) over 100 posterior predictive data replicates for diatom (top) and dinoflagellate (bottom) log-biomasses under the Time-Space-SST biomass model. The vertical line represents the observed value of the test statistic The Bayesian p-values displayed in each panel are all in the range 0.25-0.75, far from the extremes 0 and 1, implying that the model predictions do not systematically deviate from the data.
